# Draft Genome of the Korean smelt *Hypomesus nipponensis* and its transcriptomic responses to heat stress in the liver and muscle

**DOI:** 10.1101/2021.03.26.437215

**Authors:** Biao Xuan, Jongbin Park, Sukjung Choi, Inhwan You, Bo-Hye Nam, Eun Soo Noh, Eun Mi Kim, Mi-Young Song, Younhee Shin, Ji-Hyeon Jeon, Eun Bae Kim

## Abstract

Pond smelt (*Hypomesus nipponensis*) is a cold-freshwater fish species as a winter economic resource of aquaculture in South Korea. Due to its high susceptibility to abnormal water temperature from global warming, a large number of smelt die in hot summer. Here, we present the first draft genome of *H. nipponensis* and transcriptomic changes in molecular mechanisms or intracellular responses under heat stress. We combined Illumina and PacBio sequencing technologies to generate the draft genome of *H. nipponensis*. Based on the reference genome, we conducted transcriptome analysis of liver and muscle tissues under normal (NT, 5°**C**) versus warm (HT, 23°**C**) conditions, to identify heat stress-induced genes and gene categories. We observed a total of 1,987 contigs, with N_50_ of 0.46 Mbp with a largest contig (3.03 Mbp) in the assembled genome. A total number of 20,644 protein coding genes were predicted, and 19,224 genes were functionally annotated: 15,955 genes for Gene Ontology (GO) terms; and 11,560 genes for KEGG Orthology (KO). We conducted the lost and gained genes analysis compared with three species that human, zebrafish and salmon. In the lost genes analysis, we detected smelt lost 4,461 (22.16%), 2,825 (10.62%), and 1,499 (3.09%) genes compare with above three species, respectively. In the gained genes analysis, we observed smelt gain 1,133 (5.49%), 1,670 (8.09%), and 229 (1.11%) genes compare with above species, respectively. From transcriptome analysis, a total of 297 and 331 differentially expressed genes (DEGs) with False discovery rate (FDR) < 0.05 were identified in the liver and muscle tissues, respectively. Gene enrichment analysis of DEGs indicates that up-regulated genes were significantly enriched for lipid biosynthetic process (GO:0008610, P < 0.001) and regulation of apoptotic process (GO:0042981, P < 0.01), and down-regulated genes by immune responses such as myeloid cell differentiation (GO:0030099, P < 0.001) in the liver under heat stress. In muscle tissue, up-regulated genes were enriched for hypoxia (GO:0001666, P < 0.05), transcription regulator activity (GO:0140110, P < 0.001) and calcium-release channel activity (GO:0015278, P < 0.01), and down-regulated genes for nicotinamide nucleotide biosynthetic process (GO:0019359, P < 0.01). The results of KEGG pathway analysis were similar to that of gene enrichment analysis. The draft genome and transcriptomic of *H. nipponensis* will be used as a useful genetic resource for functional and evolutionary studies. Our findings will improve understanding of the molecular mechanisms and heat responses and will be useful for predicting survival of the smelt and its closely related species under global warming.

## Introduction

Pond smelt (*Hypomesus nipponensis*) is a member of the family Osmeridae native to several countries, such as South Korea, China, Japan and the United States. Smelt as a cold-freshwater fish species it is called “Bing eo” in Korea, which means ice-fish, because it resides in low-temperature water that 0 to 15°C, and it is very popular as winter fishing and food in Korea, so there are many smelt festivals are held in winter. Smelt is an anadromous species(Saruwatari, Lopez, & Pietsch, 1997), however, now it is cultivated in many reservoirs in South Korea. In a previous study, smelt samples from South Korea were identified as *H. nipponensis* (Choi & Kim, 2019).

The abnormal increase of temperature leads to the death of a large number of cold-water fish species. Therefore, many studies have focused on the acute and chronic heat stress of fish(Logan & Buckley, 2015). In Korea, there are local news report that the death of smelt in hot summer brings serious harm to the smelt industry. Therefore, understanding the characteristics of smelt will greatly improve the problems faced, however, there is no research on the genome or transcriptome of *H. nipponensis*. Furthermore, transcriptome analysis provides useful insight into the specific and general response to water temperature, such as heat shock response(Narum & Campbell, 2015). In addition, the cellular response of different smelt species to high water temperature is different. In longfin smelt, many DEGs associated with heat shock proteins and chaperones that up-regulated generally in fish response to heat stress. Conversely, in Delta smelt, DEGs associated with protein synthesis and metabolic processes(Basu et al., 2002). Cells generally regulate their own genes to survive under stress; however, occasionally they start programming cell death to induce apoptosis(Fulda, Gorman, Hori, & Samali, 2010). Increasing water temperature has many effects on fish such as oxidative stress, endoplasmic reticulum (ER) stress, decreased immune function, hypoxia, and reproductive dysfunction affecting egg production and fertility(Lu et al., 2016; Olsvik, Vikeså, Lie, & Hevrøy, 2013; Qiang et al., 2017). Since specific responses of different tissues were different under heat stress, many heat stress-related studies used different tissues of fish such as the liver, muscle, heart, kidney, brain, and gill (Logan & Buckley, 2015). For example, ER stress was identified in Atlantic salmon liver under heat stress, while ER stress promoted cell repair and reduced unfolded proteins; however, excessive ER stress led to cell apoptosis (Shi et al., 2019). ER stress also occurs in heat stress-treated rainbow trout *Oncorhynchus mykiss*, and it is accompanied by changes in the immune system and post-transcriptional regulation of spliceosome (Huang, Li, Liu, Kang, & Wang, 2018). In teleost coho salmon (*Oncorhynchus kisutch*) liver, severe heat stressors can affect the redox state and induce oxidative stress (Nakano et al., 2014). Moreover, water temperature affects the concentration of dissolved oxygen in water, low-oxygen concentration in water is fatal to fish, leading to hypoxia, dissolved oxygen plays an important role in maintaining biochemical and physiological processes. Hypoxia may limit energy processes and adversely affect growth, reproduction and survival(Kelly et al., 2020). There are many factors affecting the growth and survival of fish, therefore, it is necessary to clarify the mechanism of heat stress responses of smelt to reduce the mortality effectively.

In the present study, we used sequencing technology to analyze the whole genome of *H. nipponensis* and transcriptome analysis to compare gene expression to understand the intracellular response mechanism of *H. nipponensis* at different temperatures in the liver and muscle tissues.

## Materials and Methods

### Ethics statement

All the procedures performed on animals were approved by the Institutional Animal Care and Use Committee (IACUC accept number: KW-181109-2) at Kangwon National University.

### Source of H. nipponensis and genome sequencing

Whole genome sequencing was conducted for 3 samples of smelt from Inje narincheon (South Korea, 38° 04’ 34.0” N; 128° 11’ 17.2” E) and the genomic DNA was extracted from the whole bodies using DNeasy^®^ Blood & Tissue kit (Qiagen, GmbH, Hilden, Germany). Illumina libraries for whole bodies were constructed using the TruSeq Nano Sample Prep kit (Illumina, San Diego, CA, USA) following the manufacturer’s instructions. PacBio Sequel platform library was constructed using the PacBio Template Prep kit. PacBio generated reads were used for *de novo* assembly and Illumina paired-end reads were mapped to the draft genome assembly to error correct by pilon v.1.23(Walker et al., 2014). *de novo* assembly with PacBio reads was conducted by FALCON-UNZIP software 1.2.5 version, using the parameters length_cutoff = 13kb and length_cutoff_pr = 10kb. *H. nipponensis* genome size was estimated using k-mer (17-, 19- and 21-mers) analysis with Jellyfish 2.1.3 software. The assembly quality was assessed using Benchmarking Universal Single Copy Orthologs (BUSCO) v.3.0(Simão, Waterhouse, Ioannidis, Kriventseva, & Zdobnov, 2015). Figure S1 shows the analysis workflow of fish genome.

### PacBio (Iso-Seq) sequencing

Total RNA was isolated from liver and muscle tissues. For PacBio Iso-Seq data, we randomly selected 12 samples from the transcriptome analysis group (six liver and six muscle samples). The total RNA of the 12 samples were pooled for sequencing. The Iso-Seq library was prepared with SMATer PCR cDNA Synthesis kit. We used Trinity (Haas et al., 2013) software to perform a genome-guided assembly and combine Iso-Seq data to predict gene model.

### H. nipponensis genome functional annotation

The predicted genes were subjected to search against the National Center for Biotechnology Information (NCBI) non-redundant protein database (20190306 ver.) using BLASTx, with an e-value cutoff of 1E^-3^. Gene name and description were assigned to each gene based on the highest hit with BLASTx. The genes were submitted to Gene Ontology (GO) and Kyoto Encyclopedia of Genes and Genomes (KEGG) databases to obtain the functional category and pathway information using Blast2GO and KAAS (Moriya, Itoh, Okuda, Yoshizawa, & Kanehisa, 2007), respectively.

### Temperature trial and tissue sampling for RNA sequencing

Smelt were caught in the Hwacheon Dam (South Korea, 38° 07’ 01.1” N; 127° 46’ 43.1” E) and one hundred fish were randomly placed in two 20-L tanks (50 fish per tank, NT and HT groups). The fish were fed, and acclimatization occurred for 2 h at a water (from Hwacheon Dam) temperature of 5°C. After acclimation, to simulate temperature condition in natural environment, one tank was heated from 5°C to 23°C at a constant rate of 4.5°C per h by hotrod, while the other tank was maintained at the same temperature (5°C) with ice pack. Oxygen was supplied throughout the experiment. Sampling began until loss of equilibrium fish population appeared in the HT group (23°C); however, no loss of equilibrium fish population was observed in the NT group (5°C). A total of eight female fish were sampled (four active and four inanimate from NT and HT groups, respectively) and a total of twelve male fish were sampled (six active and six inanimate from NT and HT groups, respectively). After being sacrificed, muscle and liver tissues were immediately frozen in liquid nitrogen for gene expression profiling analysis.

### Constructing mRNA libraries and sequencing

Tissue mRNA was extracted from the liver and muscle samples using TRIzol^®^ reagent (Thermo Fisher Scientific, Waltham, MA, USA) and purified to remove DNA contamination using the TURBO DNA-free™ kit (Invitrogen, Carlsbad, CA, USA). RNA concentration and quality were determined using an Epoch Microplate Spectrophotometer (BioTek, Winooski, VT, USA). The thirty-seven sequencing libraries were created by reverse-transcription from 2 µg of RNA from each sample using the TruSeq Stranded mRNA Library Prep kit (Illumina, San Diego, CA, USA). Index adapters were added to identify sequences for each sample in the final data. Subsequently, the thirty-seven libraries were subjected to paired-end (2 × 101 bp) sequencing on the Illumina NovaSeq6000 Sequencing system.

### Detection of lost and gained genes

To detect lost and gained genes in the Korean smelt genome, we compared their fragmented sequences (raw reads, assembled DNA contigs and assembled RNA transcripts) with several reference genome of that the human (*Homo sapiens*), zebrafish (*Danio rerio*) and Atlantic salmon (*Salmo salar*). Sequence alignment was conducted by GASSST (v1.28) software. Figure S2 shows the method that sequence alignment coverage (%) was calculated for each gene from the alignment. As a low alignment coverage (less than 10%) indicates absence of a gene in the genome.

### Data analysis of mRNA

After removing the reads containing adaptor contamination using Trimmomatic v0.36, the clean reads were mapped to the assembled reference *H. nipponensis* genome using Bowtie v2.2.3. RSEM v1.2.31 was used to quantify gene abundances according to each *H. nipponensis* gene. Analysis of genes differentially expressed in the NT and HT groups was performed using the edgeR package (Robinson, McCarthy, & Smyth, 2009), and FDR < 0.05 was considered as a threshold to identify significantly differentially expressed genes (DEGs). GO enrichment analysis and KEGG statistical enrichment analysis of DEGs were performed using in-house perl scripts (parent-child method) and clusterProfiler R package (Yu, Wang, Han, & He, 2012), respectively. The significantly enriched GO terms were determined by Fisher’s exact test with a p < 0.05, and KEGG pathways were determined using FDR < 0.05.

### Data accessibility

*Hypomesus nipponensis* genome assembly was deposited in National Center for Biotechnology Information (NCBI) with accession number of SAMN16577099. RNA-Seq raw sequences were deposited in the Short Read Archive of the NCBI with accession number PRJNA672783.

## Results

### Genome sequence data and genome size estimation

Using the PacBio long-read sequencing approach, a 126× coverage (58.60 Gbp) of *H. nipponensis* genome was obtained from > 20-kb libraries. The short-read libraries (Illumina) generated an 80× coverage (36.99 Gbp) of genome was obtained from paired end sequencing (2×101bp) (Table S1). The genome size was estimated to be 464 Mb using k-mer (k=17) analysis (Table S2, Figure S3).

### Iso-Seq data generation

Isoform sequencing (PacBio) generated 0.49 Gb of data, which included 477,478 circular consensus sequences (CCS) reads and 1,023 mean of CCS read length, yielding 18,799 transcripts (Table S3), which were used to generate the assistant file for *ab initio* gene model prediction.

### De novo assembly of genome and quality assessment

Using long-read sequences from PacBio, the primary genome assembly was 631.85 Mbp in size, with 4,106 contigs (largest contig of 3.03 Mbp, N_50_ of 0.32 Mbp, and L_50_ of 477). Evaluation of the genome completeness analysis, against the 4,584 genes of *Actinopterygii*, BUSCO result shows the 4,290 genes (93.6%) were completely retrieved in the assembled genome, including 3,911 single-copy genes (85.3%) and 379 duplicated genes (8.3%). In addition, 84 fragmented genes (1.8%) and 210 missing genes (4.5%) were found (Table S4). After error correction using pilon, the final improved primary genome assembly of 1,987 contigs showed a total size of 499.59 Mbp, with an N_50_ of 0.46 Mbp, L_50_ of 300 contigs, GC content of 45.64%, and the largest contig length of 3.03 Mbp (Table S5). Table S6 and S7 show the Illumina and PacBio sequencing statistics. Table 1 shows the *H. nipponensis* draft genome statistics.

**Table 1.** *Hypomesus nipponensis* draft genome statistics.

| Index | *Hypomesus nipponensis* |
| --- | --- |
| Number of contigs | 1,987 |
| Total contig length | 498,930,205 |
| Estimated genome size | 464,167,205 |
| Total length / Estimated genome size | 107.48% |
| Minimum contig length | 19,619 |
| Maximum contig length | 3,031,274 |
| Average contig length | 251,097 |
| Contig length N50 | 464,523 |
| GC contents | 45.63% |
| BUSCO (Actinopterygii) complete | 93.58% |
| BUSCO (Vertebrata) complete | 95.32% |
| CEGMA complete | 92.74% |

### Gene prediction and functional annotation

Evidence-based *ab initio* gene predictions with Augustus predicted 20,644 protein-coding genes. Among these, 93.12% (19,224) predicted genes were annotated using the non-redundant database, while 6.88% (1,420) genes remained unannotated (Table S8). Figure 1.A shows the Blast top 10 hit species. Analysis of GO terms revealed that they were assigned to 15,955 genes on the three primary categories of ontology (biological process (BP), cellular component (CC), and molecular function (MF)). Figure 1B shows the top 10 assigned GO terms for each category. A total of 11,560 genes were annotated with seven categories on 447 pathways. Figure 1C shows the top three assigned KEGG terms for each category. Table 2 shows the *H. nipponensis* consensus gene model.

**Table 2.** *Hypomesus nipponensis* consensus gene model.

| Category | Index | Value |
| --- | --- | --- |
| **Gene** | Gene count | 20,644 |
|  | Maximum gene length | 207,358 (HYNIP00072CG0010) |
|  | Minimum gene length | 222 (HYNIP00473CG0050) |
|  | Average gene length | 12,073.93 |
|  | Total gene length | 249,254,225 |
|  | Genome coverage | 49.96% |
| **Exon** | Exon count | 193,124 |
|  | Exon count per gene | 9.35 |
|  | Average exon length | 178 |
|  | Total exon length | 34,376,477 |
|  | Genome coverage | 6.89% |
| **Intron** | Intron count | 172,480 |
|  | Average intron length | 1,245.81 |
|  | Total intron length | 214,877,748 |
|  | Genome coverage | 43.07% |

**Figure 1.**
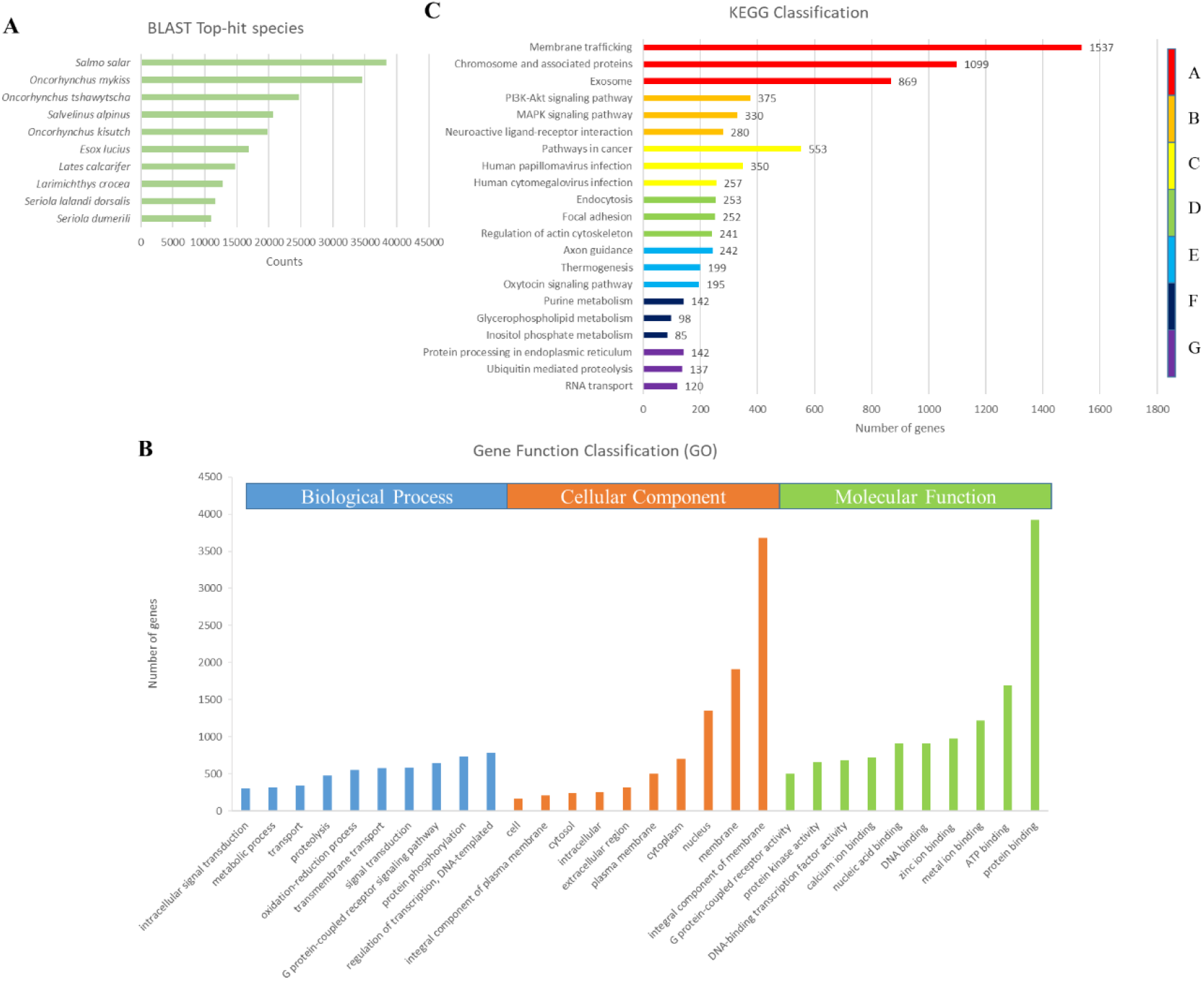
Annotation and functional classification of unigenes in genome of *H. nipponensis*. (A) Blast top 10 hit species. (B) GO analysis was performed at level 2 for the three main categories (biological process, cellular component and molecular function). (C) Pathway assignment based on the Kyoto Encyclopedia of Genes and Genomes (KEGG) database. Unigenes were classified into seven main categories (A: Genetic Information Processing; B: Metabolism; C: Organismal Systems; D: Cellular Processes; E: Human Diseases; F: Environmental Information Processing; G: Brite Hierarchies).

### Genome comparison for detection of lost and gained genes

We compared the genome of *H. nipponensis* with that of other species, including *Homo sapiens, Danio rerio*, and *Salmo salar* for detection of lost and gained genes. For lost genes found from the alignment of *H. nipponensis* genome to those of the other species, we compared 20,129, 26,612, and 48,548 genes in *H. sapiens, D. rerio*, and *S. salar*, respectively. We detected 4,461 (22.16%), 2,825 (10.62%), and 1,499 (3.09%) genes with coverage less than 10% and 6,047 (30.04%), 3,901 (14.66%), and 2,008 (4.14%) genes with coverage less than 15% (Table S9). For gained genes found from the alignment of 20,644 genes of *H. nipponensis* to those of the other species, we observed 1,133 (5.49%), 1,670 (8.09%), and 229 (1.11%) genes with coverage less than 10% and 1,911 (9.26%), 2,484 (12.03%), and 362 (1.75%) genes with coverage less than 15% (Table S10).

### GO enrichment of lost and gained genes

Gene enrichment analysis comparing smelt with human showed that smelt lost some genes involved in stimulus, reproduction, nucleotide and metal ions. Comparing with salmon showed that smelt lost chemotaxis, immune system, lipid, mitochondria, neuron, nucleotide and reproduction related genes (Table S11). In gained genes enrichment analysis, compared with human, smelt gained chemotaxis, heparin production, immune system and stimulation related genes. Compared with zebrafish and salmon, smelt gained chemotaxis and immune system related genes, respectively (Table S12).

### RNA-seq of liver and muscle transcriptome profiles at different temperature conditions

To identify differences in gene expression in smelt liver and muscle tissues at different temperatures, we constructed and sequenced twenty cDNA libraries on Illumina NovaSeq6000 platform with pair-end sequencing. The clean reads were generated by filtering the raw reads from the NT and HT groups. Table S13 and S14 shows the statistics of clean reads filtered from raw reads, and the mean quality is more than 30% for each sample.

### Gene expression profiling

A total of 297 and 331 genes were significantly differentially expressed in female liver and muscle tissues in the HT group compared to those in the NT group, respectively (|log2[Fold change]| > 1, FDR < 0.05). In the female liver tissue, 297 DEGs, including 154 up-regulated DEGs and 143 down-regulated DEGs, were identified. In the female muscle tissue, 331 DEGs, including 192 up-regulated DEGs and 139 down-regulated DEGs, were identified. In the male liver tissue, 8 DEGs, including 4 up-regulated DEGs and 4 down-regulated DEGs, were identified. In the male muscle tissue, 29 DEGs, including 9 up-regulated and 20 down-regulated DEGs, were identified (Figure 2). The volcano plot showed the DEG analysis in male liver and muscle tissues. Since the number of DEGs in male tissue is significantly less than that in female tissue, the following gene functional enrichment analysis only involves female tissue. Table S16 shows all DEGs.

**Figure 2.**
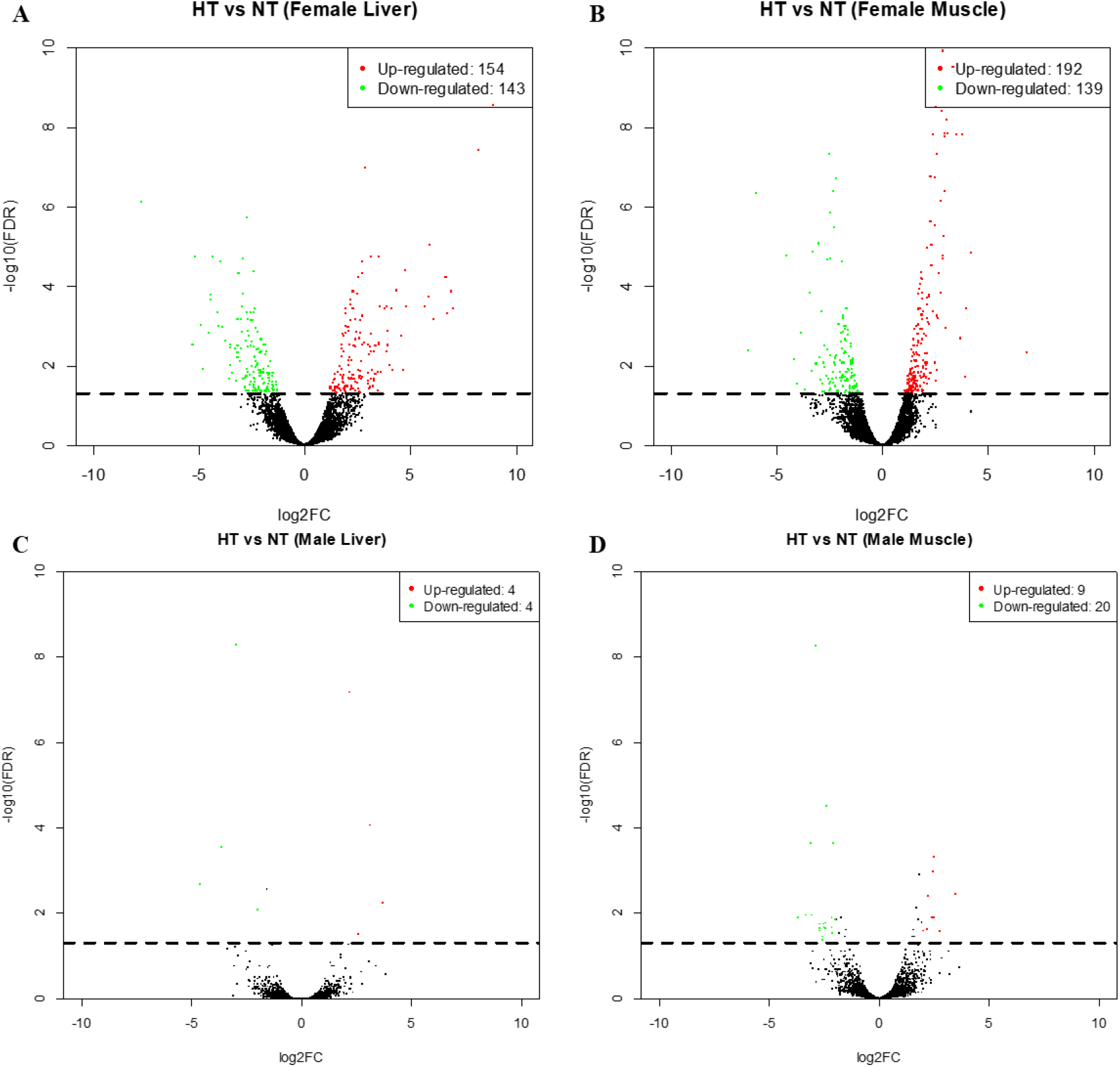
Differential expression of high-temperature in *H. nipponensis* versus low-temperature. A. DEGs in female liver. B. DEGs in female muscle. C. DEGs in male liver. D. DEGs in male muscle. The dots above the dotted line are FDR < 0.05, |log2FC|≥ 1.

### GO analysis of DEGs in the liver and muscle

In enrichment analysis, DEGs were enriched GO terms (threshold as p < 0.05 and odds ratio > 1). For the liver, 128 up-regulated and 115 down-regulated DEGs were annotated with GO terms. Up-regulated GO terms including lipid metabolism, cell death, DNA binding and stress related genes and down-regulated GO terms including immune system, membrane protein and digestion related genes (Table 3). For the muscle, 161 up-regulated and 114 down-regulated DEGs were annotated with GO terms. Up-regulated GO terms including apoptosis, hypoxia, ion transportation, transcription and component organization related genes and down-regulated GO terms including nicotinamide nucleotide and ubiquitination related genes (Table 4).

**Table 3.**
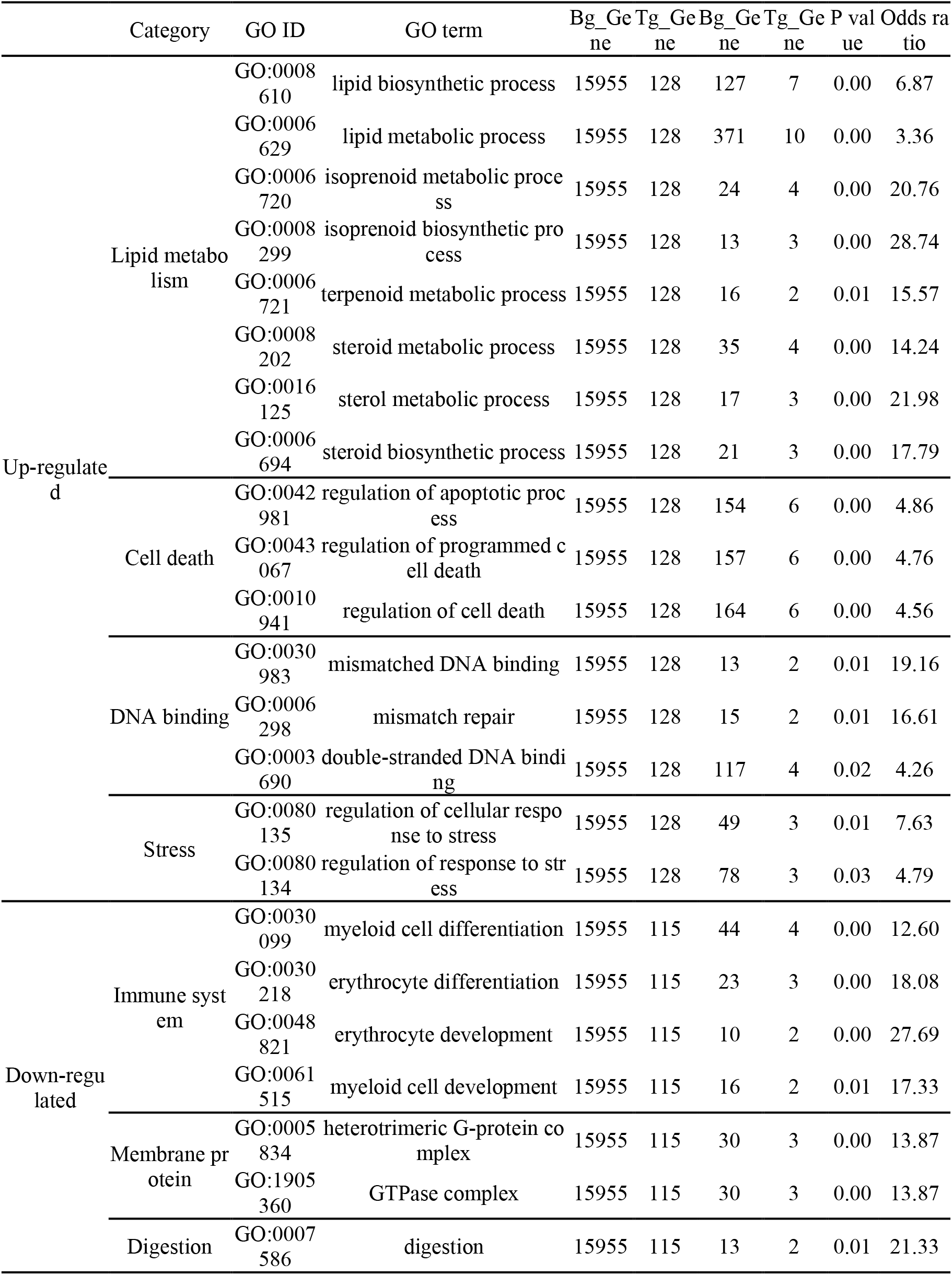
Gene enrichment analysis of DEGs in liver tissue under heat stress.

**Table 4.**
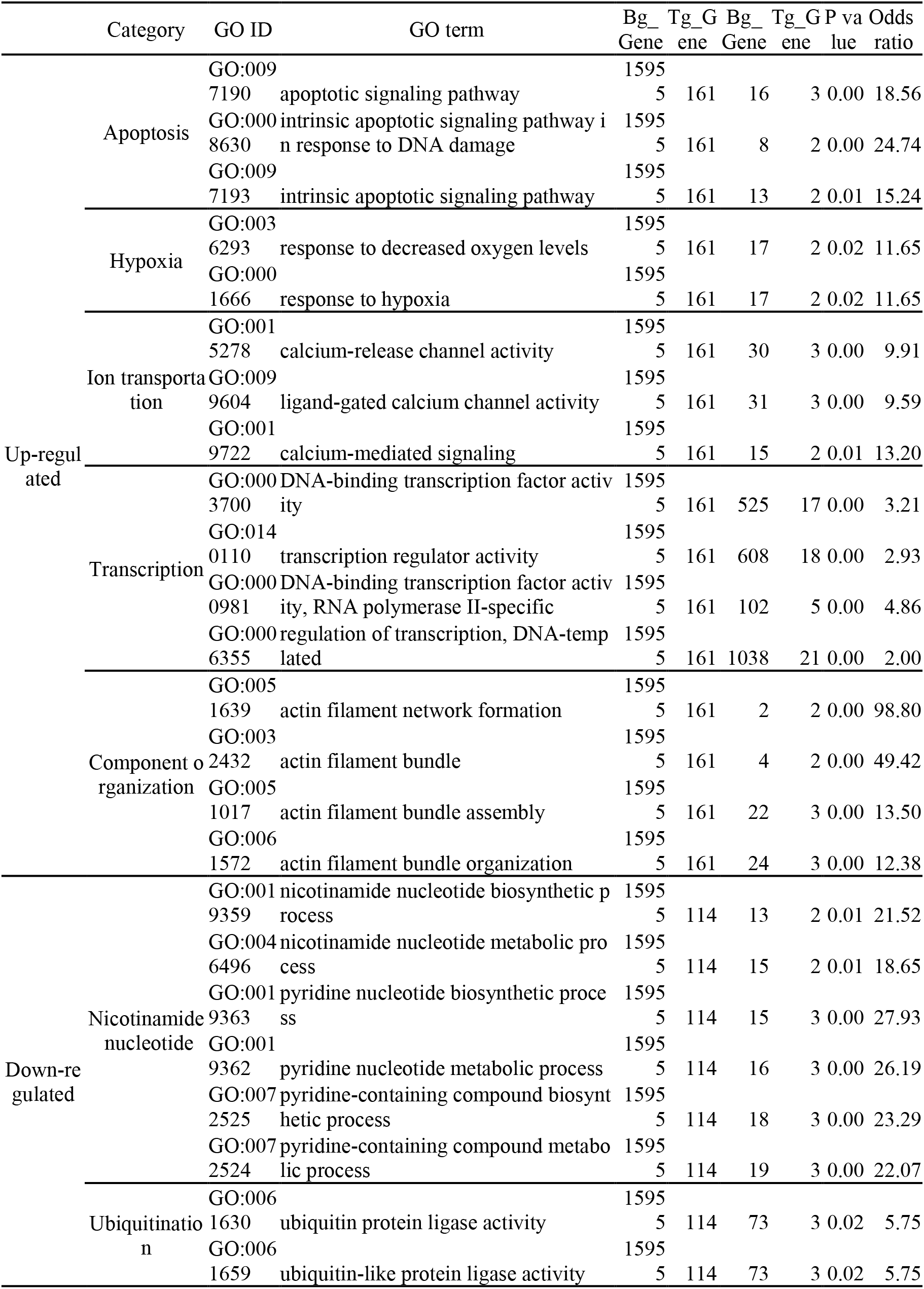
Gene enrichment analysis of DEGs in muscle tissue under heat stress.

### KEGG pathway enrichment analysis of DEGs in the liver and muscle

Four KEGG pathways were enriched in the liver tissue. One pathway (FDR < 0.1), including the terpenoid backbone biosynthesis pathway, was up-regulated, while three pathways (FDR < 0.05), including influenza A, Jak-STAT signaling, and pancreatic secretion pathways, were down-regulated. Thirty-two KEGG pathways were enriched in the muscle tissue. Mitogen-activated protein kinase (MAPK) signaling (FDR < 0.05), oxytocin signaling pathways (FDR < 0.05) and HIF-1 signaling pathway (FDR < 0.05) were up-regulated, while four pathways (FDR < 0.05), including inflammatory bowel disease (IBD), Leishmaniasis, African trypanosomiasis, and type I diabetes mellitus pathways, were down-regulated (Figure 3).

**Figure 3.**
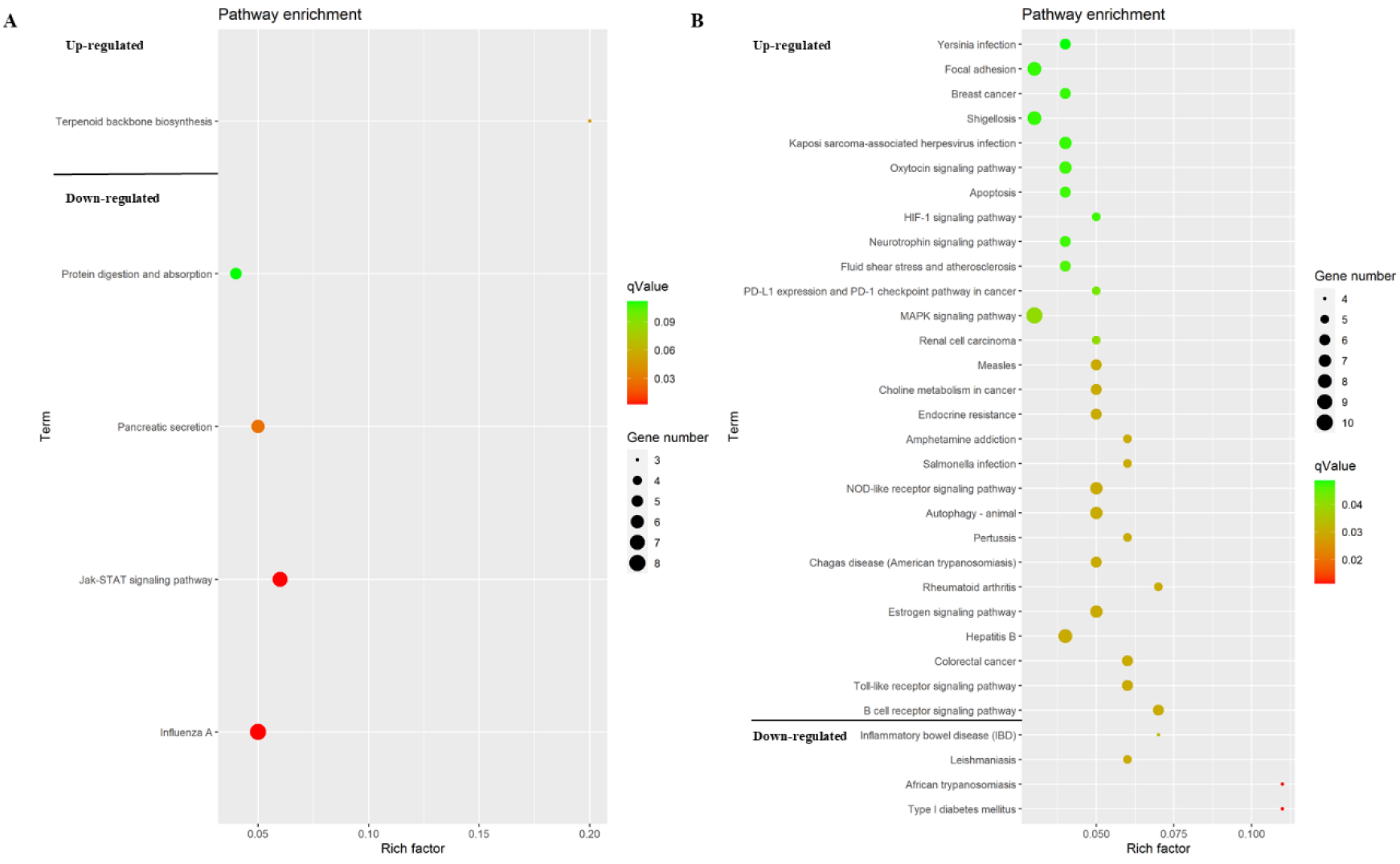
An overview of the KEGG pathway significantly enriched in DEGs in the (A) Liver, (B) Muscle. The specific pathways are plotted along the y-axis, and the x-axis indicates the enrichment factor. The size colored dots indicates the number of significantly DEGs associated with each corresponding pathway: pathway with larger-sized dots contain a higher number of genes. The color of the each dot indicates the corrected p-value for the corresponding pathway.

## Discussion

*H. nipponensis* is one of popular freshwater fish species in South Korean winter. With the rise in global warming, fish are suffering from heat stress, and a large number of deaths have been observed, resulting in great economic losses to the aquaculture industry. The present study reports the first high-quality draft genome assembly of *H. nipponensis* from the family Osmeridae (order Osmeriformes) and the analysis of transcriptome in heat stress to elucidate the mechanism of cellular response in liver and muscle tissues.

### Genome sequencing

We successfully generated *H. nipponensis* draft genome assembly of 1,987 highly contiguous contigs (498.3 Mbp), with a high N_50_ (0.47 Mbp). The estimated genome size of 464 Mbp was consistent with that of another member of Osmeridae and greater than that of *Osmerus eperlanus* (European smelt, 342.8 Mbp) in the family Osmeridae (Malmstrøm et al., 2016). Compared with other species (mammal, bird, bony fish and cartilaginous fish), the genome size of smelt is smaller than that of other species, but the GC content of genome and coding sequence (CDS) and GC content of CDS are similar to those of other species, and the number of CDS is even more than that of other species (Table S15). The genome size of *H. nipponensis* is within the range of most published fish genome size(Fan et al., 2020).

### Lost and gained genes

In the lost gene analysis, we compared *H. nipponensis* to human (*H. sapiens*), Zebrafish (*D. rerio*), and Atlantic salmon (*S. salar*). The lowest number of lost genes were observed when comparing *H. nipponensis* to *S. salar*, because salmon was a cold-water species that lived in similar water temperature. In addition, these two species are both anadromous, also, Osmeriformes are close relatives of the Salmoniformes. One of smelt species that rainbow smelt (*Osmerus mordax*) has high levels of similarity (86%) to salmonid genes(Von Schalburg et al., 2008). In the lost gene enrichment analysis, compared to human, smelt may not feel bitter taste, because smelt eat phytoplankton, they may lose their taste buds of bitterness in order to survive. In study, comparing naked mole rat (NMR) and human genome, NMR lost their receptors for bitter taste, because NMR live underground and their staple food is plant roots that are most of bitter taste, which facilitates this evolution (Kim et al., 2011). Moreover, the spermatogenesis of smelt may be different from that of human. With evolution, the genes involved in sex- and reproduction-related genes, such as mating behavior, fertilization, spermatogenesis and sex determination evolve with environmental changes(Volff, 2005). The function of cytosine is less than that of human, ammonia may not be discharged smoothly when nitrogen is excreted. Compared with salmon, sperm capacitation of smelt may be different from that of salmon. In gained genes functional enrichment analysis, compared with human, smelt have more genes related to heparin production, which may be due to the fact that living environment of smelt is low temperature, and heparin may prevent blood coagulation. Moreover, smelt may be more sensitive to light than human. Fish live in a different light environment from terrestrial species. However, water absorbs light, so as the water depth increases, the amount of light available decreases. Compared with zebrafish and salmon, the immune system of smelt is more complex.

### Transcriptome analysis

The comparative gene expression profiles in heat-stressed and normal groups are useful for understanding the mechanism of cellular responses under heat stress. In the HT group, when the temperature reached 23°C, smelt began to lose the population equilibrium, beginning to die, and there was no dead fish in the NT group. The liver is a vital metabolic organ that relates to stress response. In the gene enrichment analysis, we found that some genes involved in lipid metabolism, like lipid biosynthetic process, isoprenoid metabolic process and steroid metabolic process. Lipid is the main component of cell membrane, because acute heat stress may increase the fluidity of cell membrane, and the increase of lipid metabolism may be to maintain the stability of cell membrane. Also, lipid is the main component of fat, which can block the external heat, and lipid metabolism may increase the synthesis of fat to block the external heat. In *T. bernacchii* thermal acclimation study, there was resulted in an increase of membrane saturated fatty acids(Malekar et al., 2018). Moreover, lipid might help to maintain energy homeostasis in fish, as lipids were the basic components of sterols. Due to the membrane unique molecular structure, it has a thermal sensitive macromolecular structure. Environmental stress activates lipid metabolic enzymes and targets downstream signaling pathway(Balogh et al., 2013). Poikilothermic organisms are able to maintain their membrane fluidity for temperature fluctuation-induced cellular disturbance by regulating the composition of membrane lipids through physiological and biochemical mechanisms of homeoviscous adaptation(Mendoza, 2014). After heat shock, the plasma lipid peroxide level increase gradually and severe heat stress affects the redox state and causes oxidative stress in salmon(Nakano et al., 2014). When fish suffer from hypoxia, fat metabolism is enhanced. Isoprenoid and terpenoid metabolic process-related genes such as *HMGCS1, HMGCR*, and *FDPS* were up-regulated in liver tissue. These genes are involved in squalene synthesis. In human cancer cells study, hypoxic cells display profound accumulation of squalene(Kucharzewska, Christianson, & Belting, 2015). Additionally, it plays an anti-oxidant role that eliminates ROS produced under stress(Micera et al., 2020). Squalene synthase is up-regulated in the liver. This was the main material for the synthesis of sterols, and steroid metabolic process-related genes were up-regulated in the liver under heat stress. In other studies, heat stress led to the increase in estrogen level, and hormonal disorders led to an imbalance in the number of males and females in the population (Shi et al., 2019). Some genes related to regulation of apoptotic process were significantly up-regulated under heat stress. Although cells have different protective mechanisms under stress, the enhancement of stress can lead to cell signal interruption, extensive DNA damage, and cell apoptosis (Cheng et al., 2015). DNA damage is caused by hypoxia, leading to replication stress. Moreover, some genes related to regulation of cellular response to stress such as *PLRG1, RTEL1*, and *ING2* were up-regulated. *PLRG1* is involved in pre-mRNA splicing as a component of the spliceosome critical for heat environment adaptation (Huang et al., 2018). *RTEL1* and *ING2* play a role in DNA repair under heat stress, indicating that liver tissue has replication problems under heat stress.However, some genes related to erythrocyte characteristics, including GO term erythrocyte differentiation, erythrocyte development, and regulation of anatomical structure size, were significantly down-regulated under heat stress. Erythrocytes are cellular mediators of the immune response in teleost fish, thus, the immunity of smelt was reduced under heat stress. In addition, the genes related to cell membrane signal transduction were down-regulated in heat stress group, which indicates that it is difficult for cells to communicate with extracellular under the condition of homeostasis imbalance. Moreover, digestion related genes have also been down-regulated, suggesting that abnormal temperatures lead to feeding may be reduced, so fish should be fed less in hot summer. Study has shown that the digestive enzymes activity was significantly affected by abnormal temperature(Pimentel et al., 2015). The effect of water temperature on digestive enzymes of fish depends on species, because the optimal temperature of enzyme activity is usually in the temperature range corresponding to the fish habitat(Volkoff & Rønnestad, 2020). The muscle is greatly affected by heat stress, as, in most species, this tissue constitutes approximately 50% of the body mass. Under heat stress, apoptosis, hypoxia, ion transportation, transcription- and component organization-related genes were up-regulated. Due to high temperature, muscle cells lose calcium balance, which will cause muscle spasm, and muscle contraction-related genes that those actin-filament genes were up-regulated, also, hypoxia can cause muscle contraction. In oxygen consumption study, when the water temperature rises from 14°C to 20°C, the oxygen consumption of Delta smelt increases with the increase of temperature(Jeffries et al., 2016). In addition, nicotinamide adenine dinucleotide (NADH)- and ubiquitination-related genes associated with cell respiration and energy metabolism were down-regulated. This might be due to the decrease in collective energy metabolism caused by hypoxia.

In pathway enrichment analysis, under heat stress, terpenoid backbone biosynthesis pathway in the liver was up-regulated. The genes involved in this pathway are implicated in the final synthesis of squalene, which is an important enzyme under hypoxia condition. Immune-related pathways, including influenza A and Jak-STAT signaling pathways, were down-regulated. Digestion-related pancreatic secretion, protein digestion, and absorption pathways were down-regulated in the liver. This suggested that the digestive function of fish was disrupted under heat stress. In the muscle, several genes involved in MAPK signaling pathway were up-regulated under heat stress. A recent study has shown that MAPK signaling pathway is activated in response to ER stress (Darling & Cook, 2014). The disturbance of ER environment such as the decrease in Ca^2+^ concentration or the change in redox state can affect protein folding and processing. Once misfolded proteins accumulate, ER stress activates a series of corresponding pathways. Under heat stress, some enriched genes in oxytocin signaling pathway were also up-regulated. Oxytocin activates the signal pathways of mRNA translation during ER stress (Klein et al., 2016). HIF-1 signaling pathway was also up-regulated in the muscle tissue, indicating that smelt suffered hypoxia stress in acute increasing temperature. *HIF1A* is the main regulator of hypoxia-induced gene expression. The metabolism of smelt increases with the rise in temperature, which may lead to lack of oxygen. Smelt lives in low-temperature environments, and the oxygen solubility in low-temperature water is higher than that in high-temperature water, thus, smelt is likely to have fewer red blood cells than those of temperate fish. Hypoxic stress induced unfolded protein response (UPR) and autophagy. Hypoxia causes perturbations in the ER activity, resulting in UPR activation. These up-regulated pathways suggested that smelt suffered hypoxia and ER stress under heat stress. ER stress-related genes *NFE2L1* and *ERGIC1* were up-regulated in the muscle tissue. ER stress was also observed under heat stress in Atlantic salmon (Shi et al., 2019). IBD, Leishmaniasis, African trypaosomiasis and type I diabetes mellitus pathways were down-regulated in muscle under heat stress, these are related to immune and infectious diseases, considering that heat stress has a negative effect on the inner immune system(Lyu et al., 2018).

## Conclusions

Using long reads and short reads from PacBio Sequel and NovaSeq6000 platform, respectively, we successfully assembled the draft genome and obtained the first reference genome of *H. nipponensis*. In transcriptome analysis, smelt suffer from hypoxia and ER stress, which leads to severe oxidative stress in the body under heat stress. These results provide a better understanding of the molecular mechanisms regulating the response of *H. nipponensis* under heat stress, which will help to prevent and treat damage to fish caused by high-water temperature.

## Acknowledgments

This research was supported by a grant from the Collaborative Genome Program of the Korea Institute of Marine Science and Technology Promotion (KIMST) funded by the Ministry of Oceans and Fisheries (MOF) (No. 20180430) and the National Institute of Fisheries Scieence (R2020034). Biao Xuan was supported by the BK21 Plus Program from Ministry of Education.

